# Identification of a Highly Functional Effector CD8^+^ T Cell Program after Transplantation in Mice and Humans

**DOI:** 10.1101/2024.11.26.625263

**Authors:** Gregory S. Cohen, Joel S. Freibaum, Riley P. Leathem, Marina W. Shirkey, Robin Welsh, Sanniddhya B. Bardhan, Ryo Hatano, Chikao Morimoto, Byoung Chol Oh, Daniela Čiháková, Jonathan S. Bromberg, Scott M. Krummey

## Abstract

T cell mediated allograft rejection leads to early graft loss for kidney transplant patients. To better understand the mechanism by which T cells mediate rejection, we investigated the fate and function of graft-specific CD8^+^ T cells expressing the activated isoform of CD43 in mice and humans. Agonism of CD43 1B11 in vitro induced CD8^+^ T cell proliferation in the presence of sub-threshold antigen stimulation, and CD43 1B11 mAb treatment in vivo overcame costimulation-blockade induced tolerance to skin grafts. Relative to CD43 1B11^−^ populations, CD43 1B11^+^ CD8^+^ T cells maintained high T-bet expression along with stem-like molecules IL-7Rα and TCF-1 at both effector and memory timepoints, and were more persistent following adoptive transfer. In kidney transplant patients, graft-infiltrating CD8^+^ T cells that expressed CD43 and the glycosyltransferase GCNT1 had an effector phenotype that includes high expression of *IFNG*, *ICOS*, and perforins/granzymes. In healthy human donors and transplant candidates, the CD43 1D4 mAb clone defined antigen-experienced cytokine-producing CD8^+^ T cells. In sum, these data support a progressive differentiation model by which highly proliferative effector CD43 1B11^+^ CD8^+^ T cells infiltrate allografts also efficiently persist into memory after antigen clearance.

## 1. INTRODUCTION

T cell mediated allograft rejection is a major contributor to early graft loss.^1–3^ Multiple studies have highlighted that CD8^+^ T cells, which have potent cytolytic and cytokine-producing functions, mediate rejection in pre-clinical models and patients.^4–8^ In the widely held early precursor model of T cell differentiation, CD8^+^ T cells become either short-lived effector cell (SLEC) populations or longer-lived memory precursor populations in the initial cell divisions after antigen priming. Following viral and bacterial infection, the transcription factor T-bet is essential for CD8^+^ T cell effector function of short-lived effectors, but also sustains memory cell homeostasis.^10,16,17^ In parallel, self-renewing stem-like features of CD8^+^ T cell memory are defined by responsiveness to IL-7 and the transcription factor T cell factor 1 (TCF-1).^18–20^ While many features of effector and memory CD8^+^ T cells have been described, the allogeneic priming environment differs significantly from that of pathogen infection.^11,13,21,22^ The overarching model of CD8^+^ T cell differentiation in transplantation is not well understood.

CD43 is a transmembrane sialomucin which is broadly expressed on leukocytes, and has complex functions in cellular adhesion and trafficking, as well as cellular signaling.^23–26^ However, T cell activation elicits the core-2 O-glycosylation of CD43 by the glycosyltransferase GCNT1, ^27,28^ which is denoted by the binding of the CD43 1B11 antibody clone. Our group and others have shown that effector and memory CD8^+^ T cells upregulate expression of the activated CD43 receptor.^23,29,30^ We found that after transplantation, CD43 1B11 CD8^+^ T cells were potent effectors that expressed highly levels of Granzyme B, the cytokines IFN-γ/TNF-α, and rapidly infiltrated allografts.^29^ In the present study, we further delineated key aspects of the fate and function of this CD8^+^ T cell population in mice and humans in order to better understand the population dynamics driving T cell mediated rejection.

## 2. MATERIALS AND METHODS

### Animals

C57BL/6J, C57BL/6J Ly5.2-Cr (CD45.1), C57Bl/6-Tg(TcraTcrb)1100Mjb/J (OT-I), and C567BL/6-Tg(CAG-OVAL)916Jen/J (Act-mOVA) were obtained from The Jackson Laboratory (Bar Harbor, Maine) and used at 6-12 weeks of age. Balb/cJ, were obtained from The Jackson Laboratory and bred at Johns Hopkins University. This study was conducted in accordance with the recommendations in the Guide for the Care and Use of Laboratory Animals, and was approved by the Animal Care and Use Committee at Johns Hopkins University. Animals were housed in specific pathogen-free animal facilities at Johns Hopkins University.

### Skin transplantation and in vivo treatments

Full-thickness tail and ear skins were transplanted onto the dorsal thorax of recipient mice and secured with adhesive bandages as previously described ^31^. In some experiments, mice were treated with 250 μg/dose of CTLA-4 Ig (BioXCell) and anti-CD40L (MR1 clone, BioXCell) via i.p. injection as indicated. CD43 1B11 mAb (Biolegend) was provided at 200 μg/dose via i.p. injection.

### Heart transplantation and graft-infiltrating T cell isolation

Heterotopic heart allografts were performed as previously described via anastomosis to the abdominal aorta and pulmonary artery.^32^ To isolate graft-infiltrating cells, heart grafts were perfused with 1x PBS for 5 min, then minced and digested using Collagenase II (5000 U/mL) and DNase I (500 U/mL) (Worthington Biochemicals) in HBSS (Corning) for 30 min at 37 C with gentile agitation. Tissue was then dissociated using a GentleMACS Tissue Dissociator (Miltenyi).

### Flow cytometric cell sorting and adoptive transfer

To obtain effector CD8^+^ T cells, splenocytes and graft draining lymph nodes (inguinal, brachial, axial) from CD45.1^+^ mice grafted with Balb/c skin grafts 10-14 days prior were collected and processed to single cell suspension. For proliferation analysis, 2×10^6^ CD8^+^ T cells were isolated using CD8^+^ T cell negative selection kit (Miltenyi), and stained with CellTrace Violet (Invitrogen) prior to adoptive transfer. For fate experiments, effector CD8^+^ T cells were flow cytometrically sorted into CD44^hi^CD62L^lo^CD8^+^ T cells that were CD43 1B11^+^ and CD43 1B11^−^ using a MoFlo XDP instrument. 0.9-1.2×10^5^ purified cells were transferred via tail vein into C57BL/6J mice. Mice were grafted the following day with Balb/c skin and analyzed by flow cytometry on the indicated day post-graft as described below.

### In vitro T cell stimulation

96-well flat-bottom tissue culture plates were coated with CD3ε, CD28, or CD43 1B11 mAb (Biolegend) at the indicated concentration in 60 μL of PBS/well for 90 min at 37 C. Naïve CD8^+^ T cells isolated using CD8^+^ T Cell Isolation Kit with LS columns (Miltenyi) were stained with CellTrace Violet (Invitrogen). 2×10^5^ CD8^+^ T cells were plated per well and incubated at 37 C for 4 days in Complete R10 media comprised of RPMI 1640 with L-glutamine (Corning), 10% FBS, 100 mM HEPES, 500 uM β-mercaptoethanol, 100 U/mL penicillin/stremptomycin and analyzed by flow cytometry as described below.

### Immunofluorescence microscopy

Draining lymph nodes (dLN) were flash frozen in OCT compound (Sakura Finetek, Torrance, CA). Six μm sections were cut in triplicate using a Microm HM 550 cryostat (Thermo Fisher Scientific), fixed in cold 1:1 acetone:methanol for 5 minutes, and washed in PBS. Slides were incubated in primary antibodies and secondary antibodies diluted in PBST (PBS + 0.03% Triton-X-100 + 0.5% BSA) for 1 hour each. Slides were imaged using an Accu-Scope EXC-500 fluorescent microscope (Nikon, Tokyo, Japan). Images were analyzed with Volocity software (PerkinElmer, Waltham, MA). Mean of mean fluorescence intensity (MFI) was calculated within demarcated high endothelial venule (HEV) region. Groups were compared using quantitation of MFI multiplied by percent area (sum of HEV area with fluorescence over total HEV area x MFI).

### Murine T cell enrichment and flow cytometry

Flow cytometry and magnetic bead enrichment were performed in 1X PBS (without Ca^2+^/Mg^2+^, pH 7.2), 0.25% BSA, 2 mM EDTA, and 0.09% azide. CD45.1^+^ or L^d^ QL9^+^ CD8^+^ T cells were enriched using magnetic beads specific for fluorophores with LS columns (Miltenyi). MHC tetramers with human β2-microglobulin specific for QL9 (H-2L^d^ QLSPFPFDL) were obtained from the NIH Tetramer Core facility. Single cell suspensions were stained with Live/Dead NIR Zombie (Biolegend) and incubated with the indicated antibodies for 30-60 min at room temperature. Intracellular staining was performed using the Transcription Factor Staining Kit (eBiosciences). Samples were acquired on a Cytek Aurora (4L or 5L configuration).

### Human peripheral blood cell analysis

Healthy donor peripheral blood was collected in Vacutainer tubes containing sodium heparin (BD Biosciences) and processed according to manufacturer’s instructions to isolate mononuclear cells. For cytokine production, 1.0×10^6^ PBMC were plated in 96 well U-bottom plates in complete RPMI with L-glutamine (10% FBS, 10 mM HEPES, 10 μM β-mercaptoethanol) for 20 h at 37 C. T cells were stimulated with 0.5 μg/mL CD3ε mAb (OKT3; ThermoFisher) for 20 hrs at 37 C in the presence of 10 μg/mL GolgiStop (ThermoFisher). For flow cytometry, 0.5-1.0×10^6^ PBMC were stained with Live/Dead NIR Zombie (Biolegend), incubated with supernatant from 1D4 hybridoma for 30 min, followed by anti-mouse IgG PE, and the remaining surface antibodies as indicated. Intracellular staining was performed using the Transcription Factor Staining Kit (eBiosciences) with TNF-α and IFN-γ antibodies. Samples were acquired on a Cytek Aurora (4L and 5L) instrument.

### Human scRNAseq kidney transplant biopsy analysis

Biopsies were collected from kidney transplant patients experiencing acute rejection as described.^33^ Gene count matrices for the samples were generated by CellRanger count, and analyzed in R (4.3.0) using Seurat (4.3.0.1). Cells with excluded with fewer than 200, more than 2500 gene counts, or greater than 20% mitochondrial DNA. Analysis was filtered on mature CD8^+^ T cells defined as cells expressing at least one chain of CD3, CD8A, and not expressing CD4. The samples were integrated using the top 2000 genes and 50 dimensions. Dimensionality reductions and clustering analysis were generated using the first 12 dimensions and a resolution of 0.5. Relevant figures were generated with ggplot2 (3.4.3) and pheatmap (1.0.12).

### Quantification and Statistical Analysis

#### Statistical Analysis

All data points represent individual mice, and where individual data points are not depicted the value of n is provided in the corresponding figure legend. For analysis of absolute numbers and expression levels, paired or unpaired Student’s t-tests (two-tailed) were performed between two groups; one-way or two-way ANOVA with multiple comparison tests were used to compare multiple groups. Log-rank (Mantel-cox) test was used to evaluate graft survival between groups. Error bars represent standard error measurements (SEM). Statistics were performed using GraphPad Prism 9 or 10. Significance was determined as *p<0.05, **p<0.01, ***p<0.001, ****p<0.0001.

#### Study Approval

Healthy donors and kidney transplant candidates were isolated following protocols approved by the Johns Hopkins Institutional Review Board.

## 3. RESULTS

### 3.1 CD43 1B11 agonism with mAb provides costimulatory signal to CD8^+^ T cells

Published studies with multiple experimental systems and reagents have demonstrated that the CD43 receptor can provide either costimulatory or inhibitory signals to T cells.^25,34,35^ We stimulated purified CD8^+^ T cells with a range of plate-bound CD3ε and CD28 mAb in the presence or absence of plate-bound CD43 1B11 mAb **(Figure 1A)**. We found that a concentration of 10 μg/mL CD3ε/CD28 elicited uniform proliferation of CD8^+^ T cells **(Figure 1A)**, while 0.625 μg/mL of CD3ε/CD28 was below a threshold necessary to induce CD8^+^ T cell proliferation **(Figure 1B)**. However, the addition of CD43 1B11 mAb agonism elicited robust proliferation of CD8^+^ T cells **(Figure 1B)**. These data demonstrate that the CD43 1B11 pathway can provide a costimulatory signal to CD8^+^ T cells when a sub-threshold strength of antigen stimulation is present.

**Figure 1.**
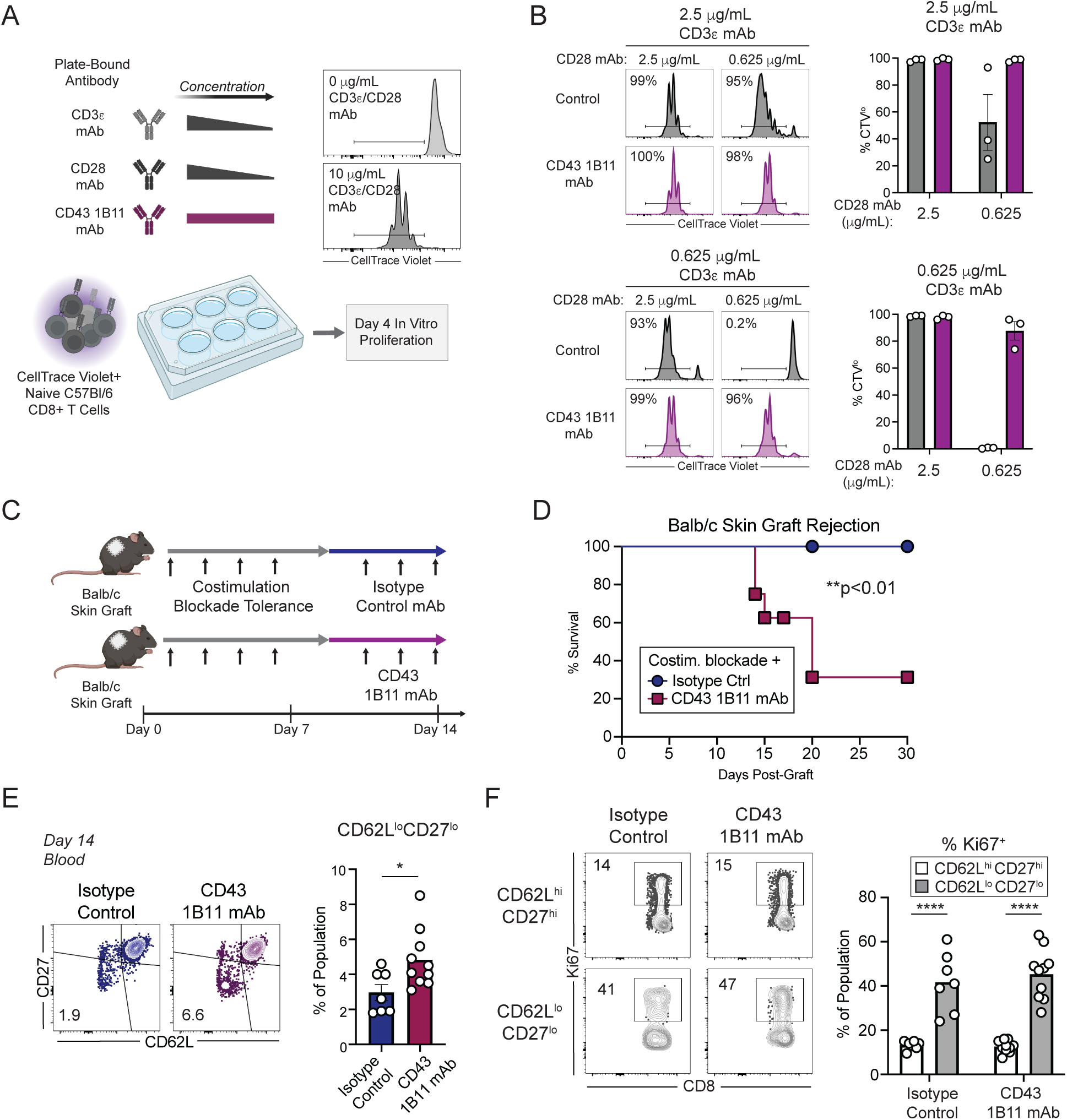
Treatment with CD43 1B11 mAb can break costimulation-blockade induced tolerance. (A) C57Bl/6 mice were provided with Balb/c skin grafts and treated with tolerance-inducing costimulation-blockade with CTLA-4 Ig and CD154 mAb (250 μg/dose, post-graft day 0, 2, 4, 6). Groups of mice were treated with IgG2a isotype control or CD43 1B11 mAb (200 μg/dose) on post-graft day 8, 11, and 14. (B) On post-graft day 14, the phenotype of CD8^+^ T cells was assessed in the blood of skin grafted mice. (C) The expression of Ki67 was assessed on populations of CD8^+^ T cells in the blood from B. (D) Skin graft survival was assessed on mice treated as described in A. Each data point depicts an individual mouse. Summary data depicts pooled results from 2 independent experiments (n=6-8 mice/group) and depicts mean ± SEM. Statistics performed by (B) Student’s t-test (2-tailed), (C) (C) 1-way ANOVA or (E) mixed-effects model (REML) with Tukey’s multiple comparisons test or (E) unpaired Student’s t-test. Error bars depict SEM. **p<0.01, ***p<0.001, ****p<0.0001.

### 3.2 CD43 mAb treatment overcomes tolerance to induce graft rejection

We reasoned that because CD43 1B11 can provide activating signals to CD8^+^ T cells and CD43 1B11^+^ CD8^+^ T cells are potent effectors, that signaling through this receptor could elicit graft rejection. We induced tolerance with combined CD28/CD40 costimulation blockade (CTLA-4 Ig and CD40L mAb; **Figure 1C**), and treated mice with CD43 1B11 mAb on day 8-14, which corresponds to the initial expression of CD43 1B11 epitope on CD8^+^ T cells ^29^. We found that while costimulation blockade induced graft survival to 30 days, treatment with CD43 1B11 elicited graft rejection in 63% of mice by day 20 **(Figure 1D)**. While costimulation blockade elicited few activated effector CD8^+^ T cells, mice treated with CD43 1B11 had a higher frequency of CD62L^lo^CD27^lo^ T cell population **(Figure 1E)**. This population is highly proliferative, as denoted by expression of Ki67 **(Figure 1F)**. Together, these data demonstrate that activation of the CD43 1B11 signaling pathway is critical for acute rejection, and can overcome transplantation tolerance.

### 3.3 The CD43^+^ICOS^+^ phenotype specifically identifies proliferative graft-specific CD8^+^ T cells

As transplantation elicits relatively weak expansion of graft-specific CD8^+^ T cells,^29,36^ we next evaluated whether the CD43 1B11^+^ICOS^+^ phenotype can be used to identify relatively rare proliferative CD8^+^ T cells after transplantation by evaluating expression of Ki67, which denotes recently divided or actively proliferating T cells in H-2^b^ C57Bl/6 mice that received an H-2^d^ Balb/c skin graft on day 14 post-grafting **(Figure 2A)**.

**Figure 2.**
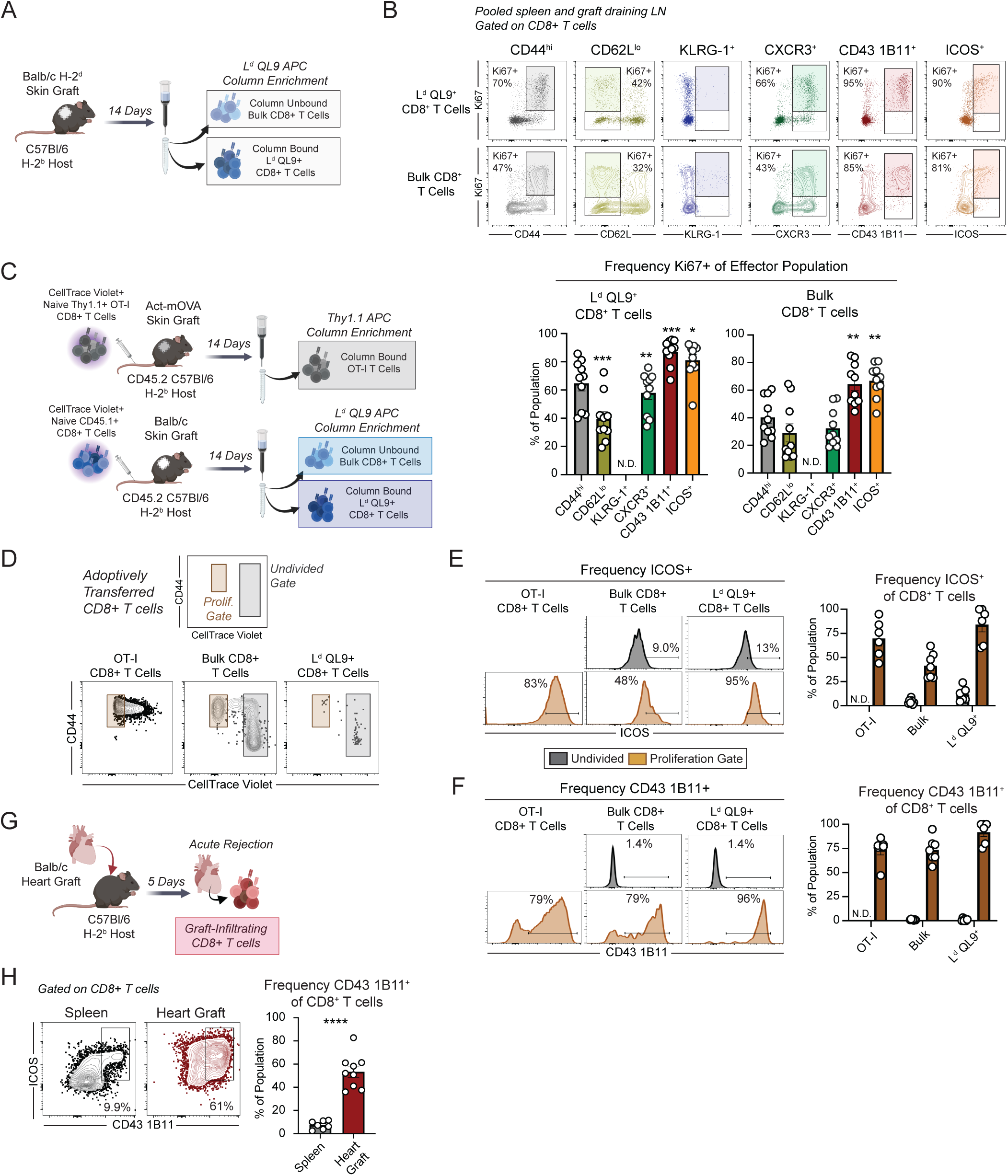
Proliferative CD8^+^ T cells are defined by CD43 1B11^+^ and ICOS^+^ phenotype after transplantation. (A) C57Bl/6 mice were provided with Balb/c skin grafts and CD8^+^ T cells were assessed on day post-graft 14 in the spleen and graft draining lymph nodes. (B) Representative flow and summary data depicting the frequency of Ki67^+^ cells among CD8^+^ T cells expressing the indicated marker. Minimal expression of KLRG-1 precluded analysis. (C) 2×10^6^ naïve CD45.1 or 1×10^6^ Thy1.1 OT-I CD8^+^ T cells labeled with CellTrace Violet were adoptively transferred into CD45.2 C57Bl/6 mice, who were provided with Balb/c or Act-mOVA skin grafts, respectively. Transferred cells were enriched using paramagnetic beads specific for APC, which stained with Thy1.1 or L^d^ QL9 tetramers. (D) Among the adoptively transferred CD8^+^ T cell populations, gates were drawn around the undivided (gray) and proliferation (orange) populations. The minimal number of undivided OT-I cells precluded further analysis. (E) The frequency of ICOS^+^ cells among each indicated adoptively transferred CD8^+^ T cell population. (F) Frequency of CD43 1B11^+^ cells among each indicated adoptively transferred CD8^+^ T cell population. (G-H) C57Bl/6 mice were provided with heterotopic Balb/c heart grafts and the phenotype of CD8^+^ T cells was assessed in the spleen and heart graft on day 5 post-graft. Each data point depicts an individual mouse. Summary data depicts pooled results from 2-5 independent experiments (n=5-10 mice/group) and depicts mean ± SEM. Statistics performed by (B) 1-way ANOVA with Dunnett’s multiple comparison test versus CD44^hi^ and (H) unpaired Student’s t-test (2-tailed). Error bars depict SEM. N.D, no data, due to low cell numbers. *p<0.05, **p<0.01, ***p<0.001, ****p<0.0001.

Among the canonical activation markers CD44^hi^ or CD62L^lo^, many CD44^hi^ or CD62L^lo^ cells were Ki67^−^, indicating that these markers are highly sensitive, but not specific, for proliferative graft-specific CD8^+^ T cells **(Figure 2B)**. As previously published, we found that very few KLRG-1^+^ cells were elicited during acute rejection **(Figure 2B)**, precluding further analysis of KLRG-1^+^Ki67^+^ cells. We found that 57.9±4.7% and 32±4.3% of CXCR3^+^ cells were Ki67^+^ in the L^d^ QL9^+^ and bulk CD8^+^ T cell compartments, respectively **(Figure 2B)**. However, both CD43 1B11^+^ and ICOS^+^ populations were highly enriched for Ki67^+^ CD8^+^ T cells **(Figure 2B)**. Among ICOS^+^, 81.1±4.1% of L^d^ QL9^+^ and 66.0±3.8% of bulk CD8^+^ T cells were Ki67^+^; among CD43 1B11^+^ cells, 87.5±2.9% L^d^ QL9^+^ and 64.5±4.7% bulk CD8^+^ T cells were Ki67^+^.

To directly visualize the proliferation of CD8^+^ T cells during acute rejection, we adoptively transferred polyclonal CellTrace Violet (CTV) labeled naïve CD45.1 CD8^+^ T cells naïve C57Bl/6 (CD45.2) hosts, followed by Balb/c skin grafting **(Figure 2C)**. In parallel, we transferred CTV labeled Thy1.1^+^ OT-I CD8^+^ T cells into C57Bl/6 hosts and provided Act-mOVA skin grafts as a comparison high-affinity antigen-specific response. After 14 days, we identified the graft-specific L^d^ QL9^+^ CD8^+^ T cells, bulk CD8^+^ T cells, or OT-I cells using magnetic column enrichment. Expression of ICOS and CD43 1B11 were largely restricted to divided cells in each population **(Figure 2D-E)**. Among divided bulk CD8^+^ T cells, the frequency of ICOS^+^ cells was 41.6±4.9%, while 73.6±4.8% were CD43 1B11^+^. L^d^ QL9^+^ CD8^+^ T cells displayed a similar profile, with divided cells being 84.2±7.5% ICOS^+^ and 91.8±4.6% CD43 1B11^+^. Thus, at this timepoint, while both ICOS and CD43 1B11 are specific for proliferating CD8^+^ T cells, CD43 1B11 is a slightly more sensitive marker for proliferating graft-specific CD8^+^ T cells.

### 3.4 CD43 1B11^+^ CD8+ T cells rapidly infiltrate heart allografts during acute rejection

We evaluated whether the functional potency of CD43 1B11^+^ CD8^+^ T cells identified in the secondary lymphoid organs corresponded to infiltration into fully vascularized heart allografts during acute rejection. In this fully allogeneic heart graft model, acute rejection occurs within 1 week post-graft ^37,38^. We evaluated the phenotype of heart graft-infiltrating CD8^+^ T cells relative to the spleen on day 5 post-graft **(Figure 2G)**, and found that while only 7.4±3.6% of CD8^+^ T cells were CD43 1B11^+^ ICOS^+^ in the spleen at this early timepoint, 53.4±14.7% of graft infiltrating CD8^+^ T cells were CD43 1B11^+^ICOS^+^ **(Figure 2H)**. These data demonstrate that CD43 1B11^+^ ICOS^+^ are a significant portion of the graft-infiltrating CD8^+^ T cells during acute rejection.

### 3.5 CD43 1B11^+^ CD8^+^ T cells maintain expression of IL-7Rα and TCF1 during acute rejection

We next evaluated the expression of key effector and stem-like markers of graft-specific CD44^hi^ CD8^+^ T cells at effector and memory timepoints **(Figure 3A)**. CD43 1B11^+^ CD8^+^ T cells were nearly universally positive for Ki67 **(Figure 3B-C).** However, CD43 1B11^+^ CD8^+^ T cells also expressed similarly high levels of IL-7Rα to naïve, CD62L^hi^ and CD62L^lo^ effector populations **(Figure 3B-C)**. CD43 1B11^+^ effectors expressed significantly higher levels of T-bet than either CD43 1B11^−^ effector population (85±2.9% CD43 1B11^+^ vs 48±3.1% CD62L^+^CD43 1B11^−^ and 52±5.1% CD62L^−^CD43 1B11^+^; **Figure 3B-C**). Finally, CD43 1B11^+^ CD8^+^ T cells also maintained relatively high levels of TCF-1 (69.9 ± 4.3%) similar to naïve and CD43 1B11^−^ effector CD8^+^ T cell populations (**Figure 3B-C**). A similar expression profile of Ki67, IL-7Rα, T-bet, and TCF-1 was observed for bulk CD44^hi^ CD8^+^ T cell populations **(Figure S1A-B)**.

**Figure 3.**
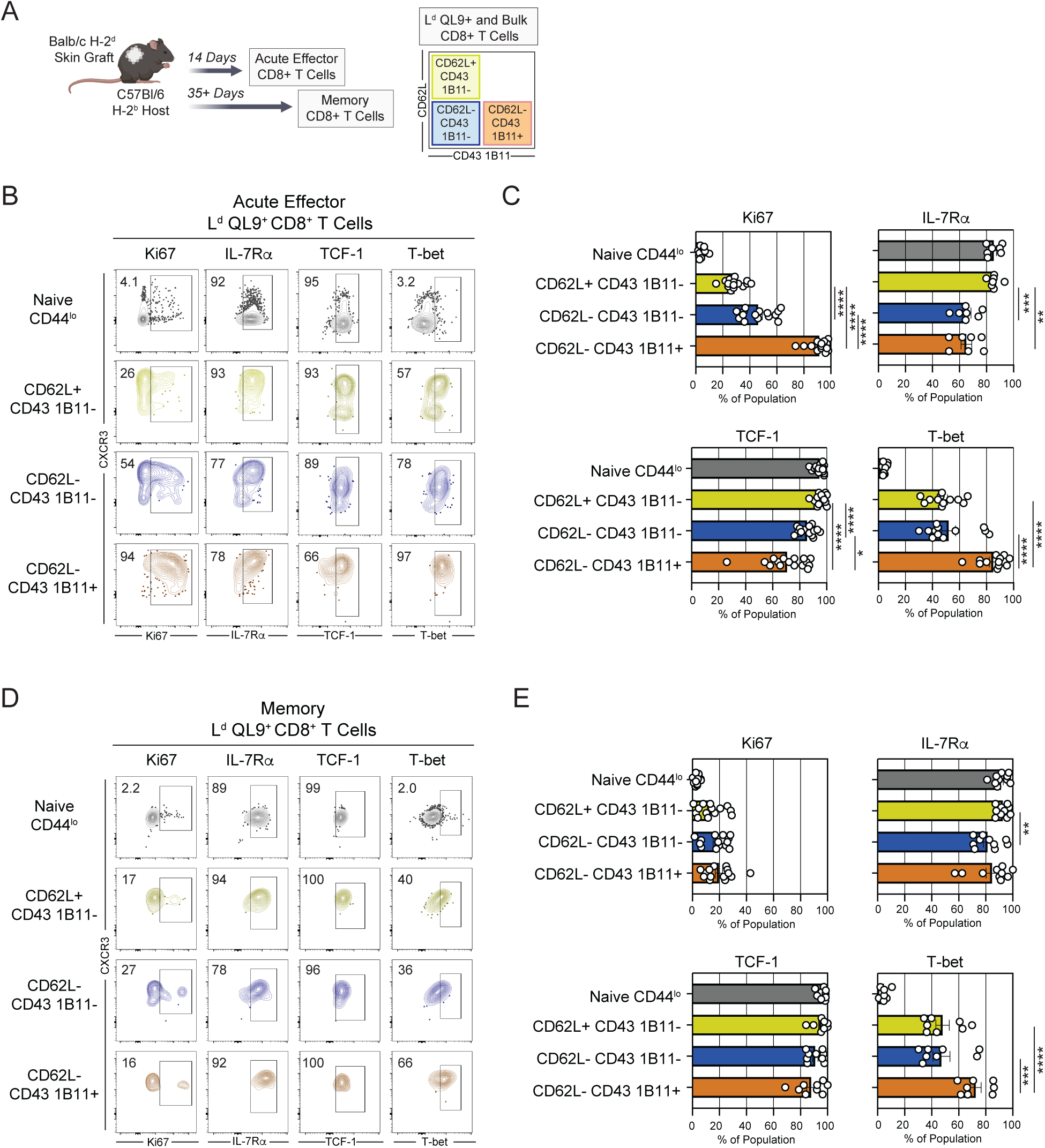
Graft-specific CD43 1B11^+^ CD8^+^ T cells have both effector and stem-like phenotypic features. (A) C57Bl/6 mice were provided with Balb/c skin grafts and CD8^+^ T cells were assessed on day 14 post-graft (Acute Effector) or day >35 post-graft (Memory) in pooled spleen and graft draining lymph nodes. Graft-specific cells were identified after enrichment using L^d^ QL9 tetramers and APC microbeads, and gated based on CD62L and CD43 1B11 expression as depicted. (B) Representative flow plots and (C) summary data depicting expression of Ki67, IL-7Rα, TCF-1, and T-bet among CD44^lo^ or CD44^hi^ L^d^ QL9^+^ CD8^+^ T cell populations at the Acute Effector timepoint. See also **Figure S1.** (D) Representative flow plots and (E) summary data depicting expression of Ki67, IL-7Rα, TCF-1, and T-bet among CD44^lo^ or CD44^hi^ L^d^ QL9^+^ CD8^+^ T cell populations at the Memory timepoint. Summary data depicts pooled results from 2-3 independent experiments (n=8-14 mice/group) and depicts mean ± SEM. Statistics performed by (C, E) 1-way ANOVA with Tukey’s multiple comparison test (excluding CD44^lo^ population used as a staining control). Error bars depict SEM. *p<0.05, **p<0.01, ***p<0.001, ***p<0.0001.

At memory timepoint (>35 days post-graft), we found that all three populations of CD8^+^ T cells were high for IL-7Rα and TCF-1 **(Figure 3D-E)**. However, CD43 1B11^+^ CD8^+^ T cells maintained higher expression of T-bet (72.6±3.9% CD43 1B11^+^ vs 48.0±5.0% CD62L^+^CD43 1B11^−^ and 47.2±6.2% CD62L^−^CD43 1B11^+^; **Figure 3D-E**). Together, these data demonstrate that CD43 1B11^+^ CD8^+^ T cells have an effector and memory program characterized by high T-bet expression versus CD43 1B11^−^ populations, but also largely maintain expression of stemness markers IL-7Rα and TCF-1.

### 3.6 Effector CD43 1B11 CD8^+^ T cells persist relative to CD43 1B11^−^ CD8^+^ T cells after transplantation

We used an adoptive transfer approach to assess the fate, and potential persistence, of CD43 1B11^+^ CD8^+^ T cells relative to CD43 1B11^−^ CD8^+^ T cells. We isolated CD62L^lo^ CD43 1B11^−^ and CD43 1B11^+^ CD8^+^ T cells from CD45.1 recipients of Balb/c skin grafts using flow cytometric sorting **(Figure 4A**). We adoptively transferred equal numbers of these cells into CD45.2 C57Bl/6 mice, challenged them with a Balb/c skin graft, and evaluated the number and phenotype of transferred populations on day 21 post-graft.

**Figure 4.**
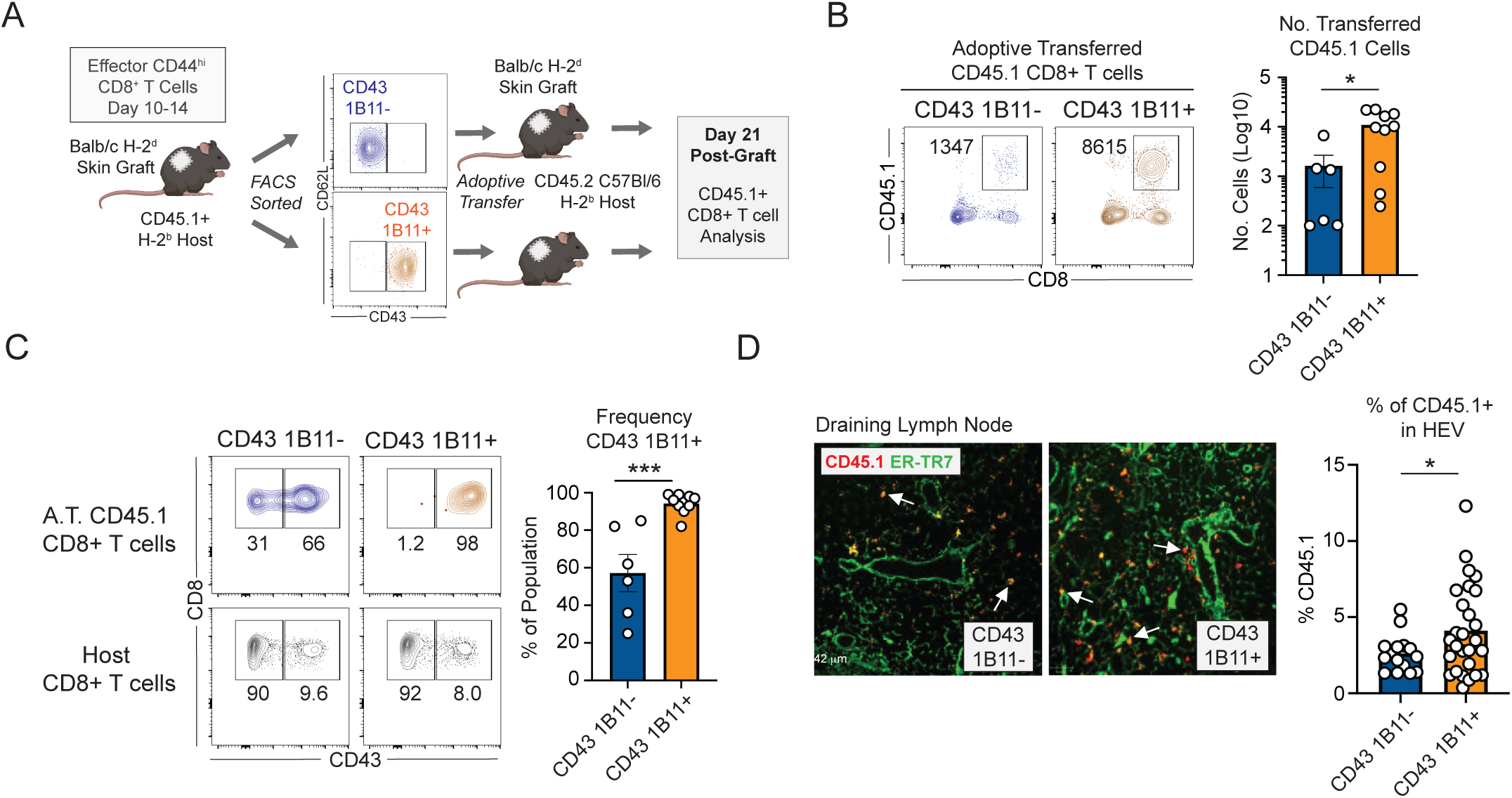
CD43 1B11^+^ CD8^+^ T cells are persistent after grafting relative to CD43 1B11^−^ CD8^+^ T cells. (A) Effector CD45.1 CD8^+^ T cells (post-graft day 10-14 after Balb/c skin grafts) were flow cytometrically sorted into CD43 1B11^+^ and CD43 1B11^−^ populations. 1×10^5^ were adoptively transferred into C57Bl/6 (CD45.2) hosts, who were provided with Balb/c skin grafts the following day. Transferred cells were assessed on post-graft day 14. (B) Recovery of CD43 1B11^+^ and CD43 1B11^−^ CD8^+^ T cells in the secondary lymphoid organs. (C) Frequency of CD43 1B11^+^ CD8^+^ T cells among adoptively transferred populations. (D) Immunofluorescence histology of draining lymph nodes depicting adoptively transferred CD45.1 CD8^+^ T cells (red) and the HEV marker ER-TR7 (green). Arrows depict localization of adoptively transferred CD45.1 CD8^+^ T cells outside (CD43 1B11^−^) or within (CD43 1B11^+^) HEV. Summary data depicts pooled results from 3 independent experiments (n=6-10 mice/group) and depicts mean ± SEM. Statistics performed by unpaired Student’s t-test (2-tailed). Error bars depict SEM. HEV, high endothelial venule. *p<0.05, **p<0.01, ***p<0.001, ***p<0.0001.

We found that relative to CD43 1B11^−^ T cells, greater numbers of CD43 1B11**^+^** T cells were recovered from secondary lymphoid tissue **(Figure 4B)**. While CD43 1B11^+^ T cells remained CD43 1B11^+^ after skin grafting, a fraction of CD43 1B11^−^ T cells converted to CD43 1B11^+^ T cells following the second skin graft challenge **(Figure 4C)**. Evaluation of the localization of these transferred populations in the draining lymph node revealed that CD43 1B11^+^ population localizes to the high endothelial venule relative to the CD43 1B11^−^ CD8^+^ T cells **(Figure 4D)**. Together, these data demonstrate that CD43 1B11^+^ CD8+ T cells appear to be a terminal phenotype, as they are formed from CD43 1B11^−^ precursors, and are persistent *in vivo*.

### 3.7 CD8^+^ T cells that are *SPN^+^GCNT1^+^* are present in the rejecting allografts of kidney transplant patients experiencing acute rejection

In order to investigate whether a population of CD43^+^ CD8^+^ T cells could be identified in the allografts of kidney transplant patients during acute rejection, we explored the single cell RNA-sequencing data generated by Shi et al ^33^ **(Figure 5A).** Among kidney transplant patients undergoing acute cellular rejection, we identified 8 clusters of CD8^+^ T cells **(Figure 5B-C)**. All clusters expressed the CD43 gene *SPN*. The because the glycosyltransferase GCNT1 is responsible for glycosylating the CD43 receptor ^13,27,28^, we reasoned that *SPN^+^GCNT1^+^*clusters could be a surrogate for CD43 1B11^+^ CD8^+^ T cells observed using flow cytometry. We found that clusters 1, 2, 4, and 6 were *SPN^+^GCNT1^+^* **(Figure 5D)**. The *SPN^+^GCNT1^+^* clusters were higher for Granzyme B (*GZB*) and lower for CD62L (*SELL*), which is consistent with the CD43 1B11^+^ CD8^+^ T cell population in mice. The *SPN^+^GCNT1^+^* clusters expressed high levels of granzymes and perforin (*GZMA, GZMH, GZMK, PRF1*). Relative to GCNT1^−^ clusters present in the allograft, they expressed elevated transcripts for IFN-γ (*IFNG*), the chemokine receptor CXCR3, and the co-receptors PD-1 and ICOS, which are markers of recent antigen encounter. Overall, these results reveal that populations of CD8^+^ T cells that are *SPN^+^GCNT1^+^*are present in rejecting kidney allografts, and these clusters have features of highly potent effectors that have recently encountered antigen.

**Figure 5.**
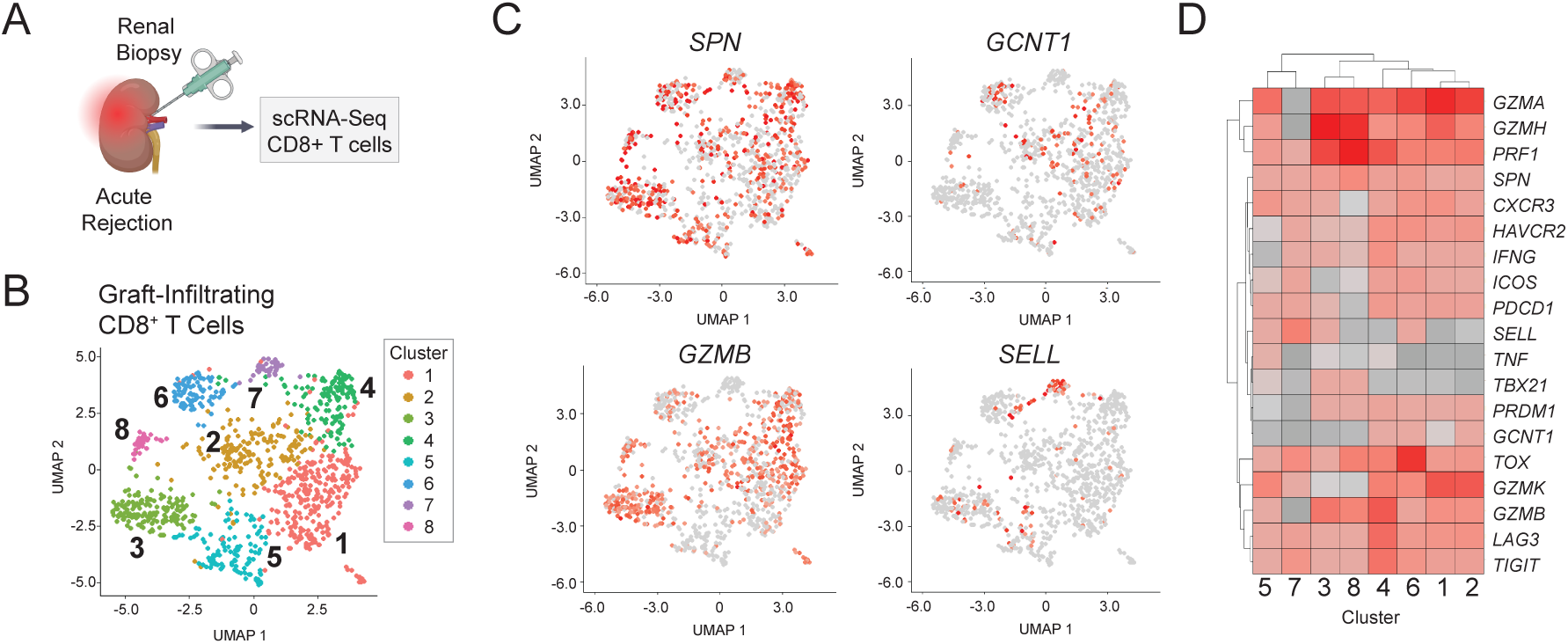
A population of effector-like graft-infiltrating CD8^+^ T cells from kidney transplant patients are *SPN^+^GCNT1^+^.* (A) Biopsies from kidney transplant patients with clinical signs of acute rejection were analyzed by single cell RNA-sequencing. (B) Hierarchical clustering and U-MAP analysis of CD8^+^ T cells from kidney allograft biopsies. (C) Expression of *SPN, GCNT1, GZMB,* and *SEL*L among CD8^+^ T cell clusters. (D) Heatmap depicting expression genes for T cell of co-receptor and effector proteins.

### 3.8 CD43 1D4 mAb identifies a population of effector CD8^+^ T in healthy human donors and kidney transplant candidates

In humans, high and low molecular weight forms of the CD43 receptor have been identified ^39^. The CD43 1D4 mAb clone identifies a large core-2 O-glycosylated isoform of CD43 on human T cells ^39^. In healthy donors, we found that a fraction of CD14^+^ monocytes and CD8^+^ T cells stained for CD43 1D4 **(Figure 6A)**. We investigated whether the CD43 1D4^+^ CD8^+^ T cells corresponded with the canonical naïve (CCR7^+^CD45RA^hi^) or antigen-experienced T cell populations: central memory (T_CM_, CCR7^+^CD45RA^lo^), effector memory (T_EM_, CCR7^−^CD45RA^lo^), or T effector memory RA^+^ (T_EMRA_, CCR7^−^CD45RA^+^) populations. While CD43 ID4^−^ CD8^+^ T cells were predominantly found within the naïve compartment, CD43 1D4^+^ CD8^+^ T cells were found predominantly within antigen-experience fractions **(Figure 6B).** Further exploration of the phenotype of these cells revealed that CD43 1D4^+^ CD8^+^ T cells had lower expression of CD27 and higher expression of CXCR3 and ICOS than CD43 1D4^+^ CD8^+^ T cells **(Figure 6C)**.

**Figure 6.**
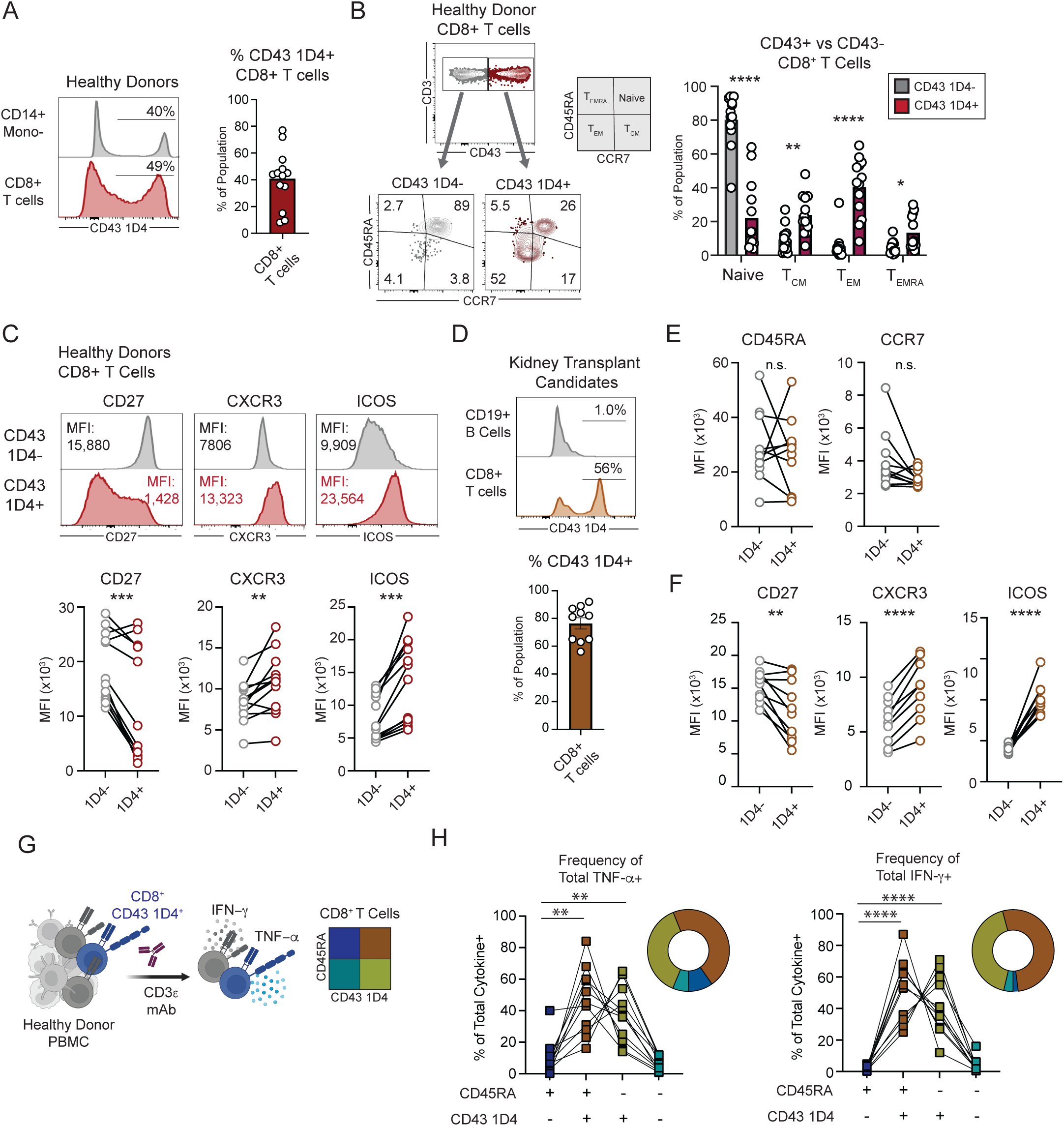
A population of human CD8^+^ T cells are CD43 1D4^+^ and have an antigen-experienced effector phenotype. (A) The frequency of peripheral healthy donor CD14^+^ monocytes and CD8^+^ T cells that express the CD43 epitope 1D4. (B-C) Expression of CD45RA and CCR7 among CD43 1D4^+^ and 1D4^−^ CD8^+^ T cells from healthy donors. (C) Representative flow plots and summary data showing expression of CD27, CXCR3, and ICOS among CD43 1D4^+^ and 1D4^−^ CD8^+^ T cell populations. (D) Frequency of CD43 1D4^+^ CD8^+^ T cells among kidney transplant candidates. (E) Expression of CD45RA, CCR7 among CD43 1D4^+^ and CD43 1D4^−^ CD8^+^ T cells from kidney transplant candidates. (F) Expression of CD27, CXCR3, and ICOS, among CD43 1D4^+^ and CD43 1D4^−^ CD8^+^ T cells from kidney transplant candidates. (G) Healthy donor PBMC were stimulated for 20-24 h with CD3ε mAb (OKT3) and assessed for TNF-α and IFN-γ production. See also **Figure S2**. (H) Summary data depicting the proportion of TNF-α (left) or IFN-γ (right) produced by each CD45RA and CD43 1D4 quadrant population. Each data point depicts an individual human donor. Summary data depicts pooled results from (A-C) 3 independent experiments (n=12-13 individuals/group), (D-F) 2 independent experiments (n=10 individuals/group), (H) 3 independent experiments (n=11 individuals/group). Summary data and depicts mean ± SEM. Statistics performed by (B) 2-way ANOVA with Sidak’s multiple comparisons test and (C, E) paired Student’s t-test (2-tailed), (G) 1-way ANOVA with Dunnett’s multiple comparisons test. Error bars depict SEM. *p<0.05, **p<0.01, ***p<0.001, ****p<0.0001.

We next investigated whether CD43 1D4 was expressed on CD8^+^ T cells from patients with end-stage renal disease who are waitlisted for kidney transplantation. Similar to healthy donors, a population of CD8^+^ T cells expressed CD43 1D4 **(Figure 6D)**. We found that relative to CD43 1D4^−^, CD43 1D4^+^ CD8^+^ T cells had variable and statistically similar expression of CD45RA and CCR7, demonstrating that this population did not correspond to a canonical T_EM_, T_CM_, or T_EMRA_ populations **(Figure 6E).** CD43 1D4^+^ CD8^+^ T cells from these patients also displayed lower expression of CD27 and higher expression of ICOS and CXCR3 **(Figure 6F).** Overall, these data demonstrate that CD43 1D4 identifies a population of antigen-experienced CD8^+^ T cells in human healthy donors and renal transplant candidates.

### 3.9 CD43 1D4^+^ CD8^+^ T cells are efficient cytokine producers relative to CD43 1D4^−^ CD8^+^ T cells

To evaluate the functional capacity of CD43 1D4^+^ and CD43 1D4^−^ CD8^+^ T cells, we stimulated PBMC from healthy donors with CD3ε mAb for 18-20 hours and evaluated CD8^+^ T cells based on CD45RA and CD43 1D4 expression **(Figure S2A and Figure 6G).** We found that CD3ε mAb stimulation increased CD43 1D4 expression among both CD45RA^hi^ and CD45RA^lo^ populations **(Figure S2B)**. This change was very modest among CD45RA^lo^ cells, and was more pronounced among CD45RA^hi^ cells, presumably due to activation of a subset of naïve CD8^+^ T cells. When we compared the cytokine production among CD43 1D4^+^ and 1D4^−^ fractions, we found that both CD45RA^hi^ and CD45RA^lo^ CD43 1D4^+^ fractions produced elevated levels of TNF-α and IFN-γ relative to the CD43 1D4^−^ populations **(Figure 6D).** In total, these data demonstrate that CD43 1D4 is a marker of antigen-experienced and acute effector CD8^+^ cells that are functional as determined by cytokine production.

## 4. DISCUSSION

In this study, we built upon previous work that showed graft-specific L^d^ QL9^+^ CD8^+^ T cells express the activated glycoform of the siaolomucin CD43, defined by the 1B11 mAb.^29^ Specifically, we sought to evaluate whether CD43 1B11 a marker of short-lived, highly proliferative CD8^+^ T cells, analogous to KLRG-1^+^ short-lived effector CD8^+^ T cells.

We found that treatment with CD43 1B11 mAb *in vivo* was able to break costimulation-blockade induced tolerance and elicit skin graft rejection in the majority of mice after Balb/c skin grafting. To our knowledge, prior studies using CD43 mAb have not distinguished between global signaling through the CD43 receptor and the specific function of activated CD43 1B11^+^ receptor.^25,34,35^ Our results demonstrate that CD43 1B11 agonism provides a costimulatory signal to CD8^+^ T cells that can overcome sub-threshold antigen signaling, and that this pathway is critical for alloimmunity *in vivo*.

We found that CD43 1B11^+^ status was highly sensitive for proliferating CD8^+^ T cells during acute rejection, and that CD43 1B11^+^ CD8^+^ T cells are a significant fraction of graft-infiltrating CD8^+^ T cells during acute rejection of heart allografts. Consistent with a highly potent effector phenotype, CD43 1B11^+^ CD8^+^ T cells had higher levels of T-bet expression versus CD43 1B11^−^ CD8^+^ T cell populations. However, in contrast to terminally differentiated effector CD8^+^ T cells, CD43 1B11^+^ CD8^+^ T cells also maintained expression of memory-precursor and stem-like makers TCF-1 and IL-7Rα, and were more persistent in adoptive transfer experiments. TCF1 is associated with the survival and proliferative capacity of T cells in a variety of settings,^18,19,43,44^ and TCF1 expression in CD4^+^ T cells corresponds with self-renewal capacity of effectors in transplantation.^43^ Further work is needed to understand the importance of the heterogeneity of TCF-1 expression within the CD43 1B11^+^ population, including whether this population is predominantly self-renewing or fed by CD43 1B11^−^ cells.

In total, these results demonstrate that CD43 1B11^+^ CD8^+^ T cells possess a combination of effector and stem-like features: potent effector capacity (cytokine production, rapid graft infiltration, localization near the HEV), robust proliferation, and enhanced survival. In contrast to the early memory-effector precursor model after viral infection, our findings point to a progressive differentiation model of CD8^+^ T cell activation in which CD43 1B11^+^ CD8^+^ T cells dominate the proliferative cytotoxic response during the acute phase, but also efficiently persist into memory after antigen clearance. Additional studies are needed to further dissect the population dynamics of the CD43 1B11^+^ population and elucidate the full transcriptional program driving its function. Future studies are needed to explore the potential approaches to therapeutically target them in mice and humans.

## ACKNOWLEDGEMENTS/FUNDING

We thank members of the Johns Hopkins Department of Pathology Division of Immunology for helpful discussions. We appreciate Hao Zhang and the Bloomberg School of Public Health Flow Cytometry and Cell Sorting Core for excellent cell sorting, the Johns Hopkins Ross Flow Cytometry Core, and the NIH Tetramer Core Facility for tetramer reagents and technical advice. We thank the Johns Hopkins Collaborative Outcomes Research in Endocrine Surgery (CORES) for human subject sample collection and storage. This work was supported by National Institute of Allergy and Infection Disease (NIAID) R56 AI179856 (S.M.K.), American Society of Transplantation Research Network Career Transition Grant (S.M.K.), KidneyCure Carl W. Gottschalk Research Scholar Grant (S.M.K.), NIAID R01 AI114496 (J.S.B.), National Heart, Lung, and Blood Institute (NLBI) R01HL118183 (D.C.), NHLBI R01HL136586 (D.C.), National Cancer Institute R01CA289729 (D.C.), The American Heart Association 23EIA1040103 (D.C.), Global Autoimmune Institute (D.C.), and the Matthew Poyner MVP Memorial Myocarditis Research Fund (D.C.).

## DATA AVAILABILITY STATEMENT

Further information and requests for resources and reagents should be directed to the lead contact, Scott Krummey

## AUTHOR CONTRIBUTIONS

Conceptualization, S.M.K., G.S.C., J.S.B., R.P.L., J.S.F., B.O., D.C.; methodology, G.S.C., J.S.F., M.W.S., R.H., S.M.K., C.M., J.S.B., R.W.; formal analysis, S.M.K., G.S.C., M.W.S.; investigation, G.S.C., J.S.F., R.P.L., M.W.S., R.H., R.W.; writing, G.S.C., S.M.K.; visualization, S.M.K., G.S.C., J.S.F., R.P.L.; supervision, S.M.K., J.S.B., C.M., D.C.

---

**Figure S1.**
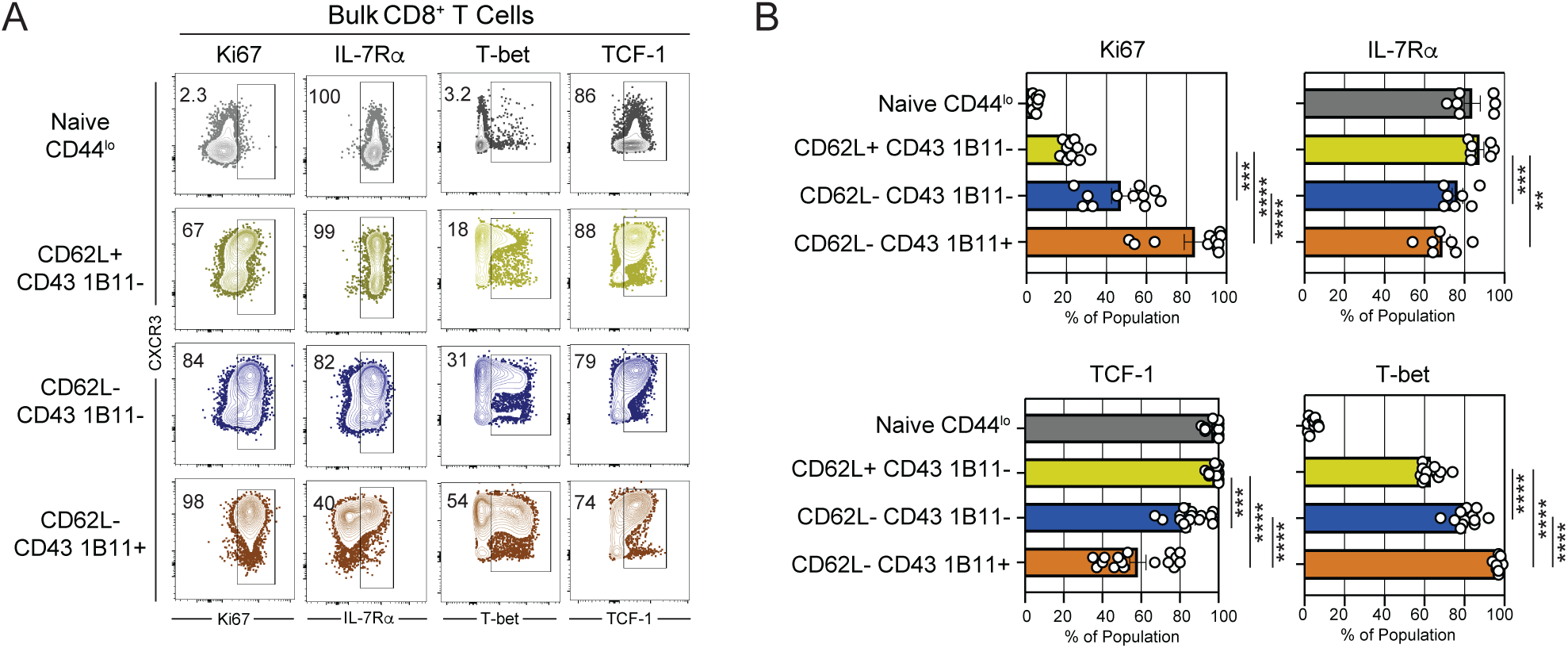
CD43 1B11^+^ CD8^+^ T cells have both effector and stem-like phenotypic features. Related to Figure 3. C57Bl/6 mice were provided with Balb/c skin grafts and CD8^+^ T cells were assessed on day 14 post-graft (Acute Effector) in the pooled spleen and graft draining lymph nodes. (A) Representative flow plots and (B) summary data depicting expression of Ki67, IL-7Rα, TCF-1, and T-bet among CD44^lo^ or CD44^hi^ L^d^ QL9^+^ CD8^+^ T cell populations at the Acute Effector timepoint. Summary data depicts pooled results from 2-3 independent experiments (n=8-14 mice/group) and depicts mean ± SEM. Statistics performed by 1-way ANOVA with Tukey’s multiple comparison test (excluding CD44^lo^ population used as staining control). Error bars depict SEM. *p<0.05, **p<0.01, ***p<0.001, ***p<0.0001.

**Figure S2.**
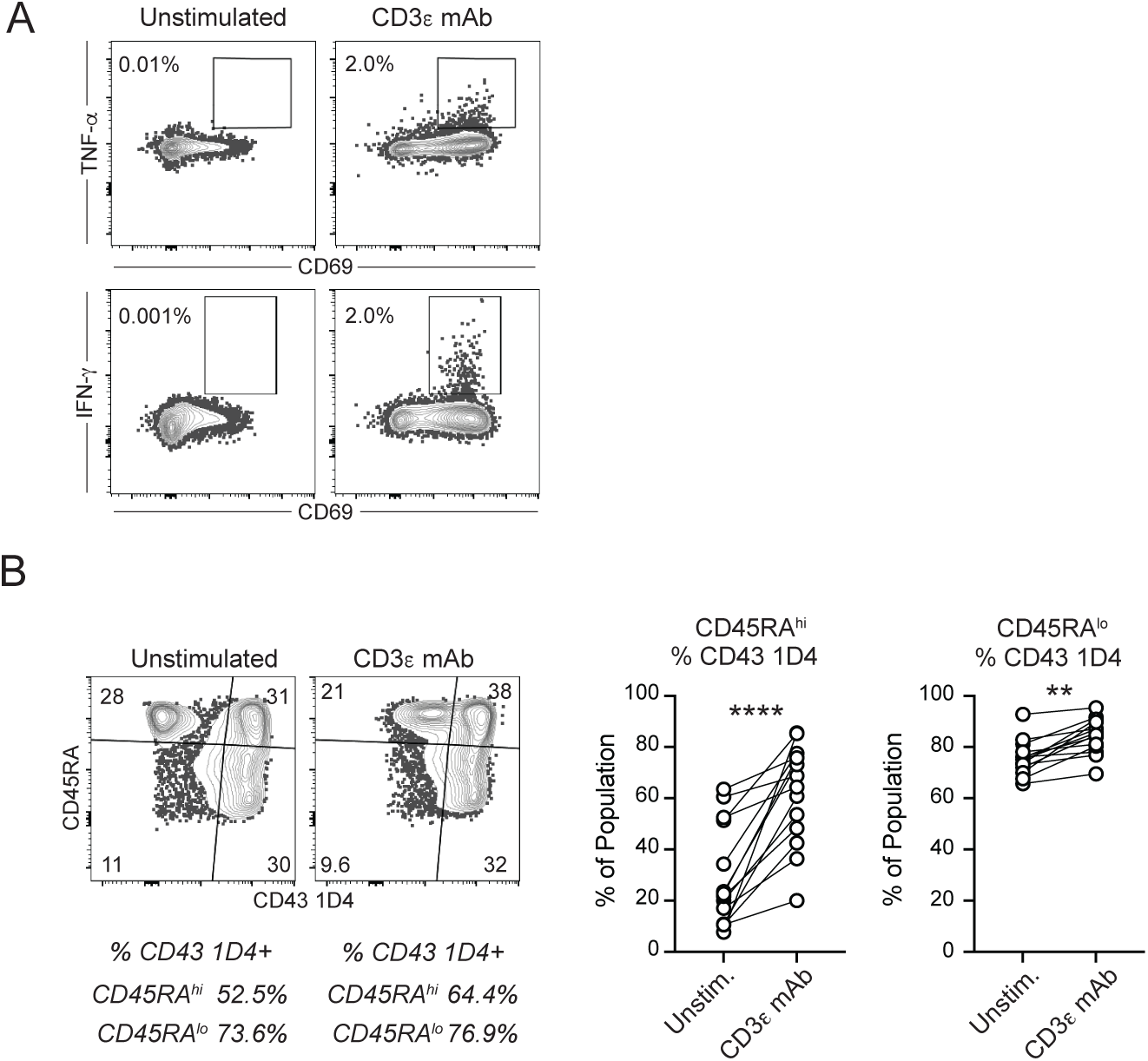
Evaluation of cytokine production by human CD43 1D4^+^ CD8^+^ T cell populations. Related to Figure 6. (A) Healthy donor PBMC were stimulated for 20-24 h with CD3ε mAb (OKT3) and assessed for TNF-α and IFN-γ production. (B) Representative flow cytometry plots and summary data showing expression of CD43 1D4 expression among CD45RA^hi^ and CD45RA^lo^ populations of CD8^+^ T cells. Summary data depicts pooled results from 3 independent experiments (n=11 individuals/group), and depict mean ± SEM. Statistics performed by (B) paired Student’s t-test (2-tailed), **p<0.01, ****p<0.0001

## SUPPLEMENTAL INFORMATION

Additional supporting information may be found online in the Supporting Information section.

